# Tracking activity patterns of a multispecies community of gymnotiform weakly electric fish in their neotropical habitat without tagging

**DOI:** 10.1101/550814

**Authors:** Jörg Henninger, Rüdiger Krahe, Fabian Sinz, Jan Benda

## Abstract

Field studies on freely behaving animals commonly require tagging and often are focused on single species. Weakly electric fish generate a species- and individual-specific electric organ discharge (EOD) and therefore provide a unique opportunity for individual tracking without tagging. We here present and test tracking algorithms based on recordings with submerged electrode arrays. Harmonic structures extracted from power spectra provide fish identity. Localization of fish based on weighted averages of their EOD amplitudes is found to be more robust than fitting a dipole model. We apply these techniques to monitor a community of three species, *Apteronotus rostratus*, *Eigenmannia humboldtii*, and *Sternopygus dariensis*, in their natural habitat in Darién, Panamá. We found consistent upstream movements after sunset followed by downstream movements in the second half of the night. Extrapolations of these movements and estimates of fish density obtained from additional transect data suggest that some fish cover at least several hundreds of meters of the stream per night. Most fish, including *Eigenmannia*, were traversing the electrode array solitarily. From *in-situ* measurements of the decay of the EOD amplitude with distance of individual animals we estimated that fish can detect conspecifics at distances of up to 2 m. Our recordings also emphasize the complexity of natural electrosensory scenes resulting from the interactions of the EODs of different species. Electrode arrays thus provide an unprecedented window into the so-far hidden nocturnal activities of multispecies communities of weakly electric fish at an unmatched level of detail.

**Summary statement:** Detailed movement patterns and complex electrosensory scenes of three species of weakly electric fish were tracked without tagging using a submerged electrode array in a small Neotropical stream.

## Introduction

Studying the sensory ecology of a species (Endler and Basolo, 1998) or the statistics of the natural sensory environment that animals perceive (Laughlin, 1981; Smith and Lewicki, 2006) is ideally based on highly resolved observations of freely behaving animals in their natural habitats (Henninger et al., 2018). However, most outdoor studies require loggers or tags to be mounted on the animals (e.g., Menzel et al., 2005; Baktoft et al., 2015; Strandburg-Peshkin et al., 2015; Cvikel et al., 2014; Flack et al., 2018). While recent advances in high-precision technologies for tracking behaving animals allow for a new level of high-throughput studies in the neurosciences (Anderson and Perona, 2014; Egnor and Branson, 2016; Gomez-Marin et al., 2014; Mathis et al., 2018), ecology and animal behavior (Dell et al., 2014; Hughey et al., 2018), tracking animals in their natural habitats remains notoriously difficult: the size and complexity of natural, cluttered environments are challenging for video-based techniques (Dell et al., 2014). Instead, many studies have applied animal-mounted senders and loggers for tracking, e.g., baboons, storks, large fish, and even insects (Strandburg-Peshkin et al., 2015; Flack et al., 2018; Baktoft et al., 2015; Menzel et al., 2005). In addition to tracking identity and movements, communication signals or physiological parameters are of interest and would usually require additional sensors to be mounted on the animal (Cvikel et al., 2014), or to be distributed in the field (Rodriguez-Munoz et al., 2010). This usually restricts observations to a subset of the animals in a population of interest and to a single species.

Notably, weakly electric fish provide a unique opportunity for monitoring identity, movement, and communication signals of individuals of multiple species without disturbing the animals: these fish continuously generate an electric field by species-specific electric organ discharges (EODs; Fig. 1 A, Kramer et al., 1981; Hopkins and Heiligenberg, 1978; Turner et al., 2007). By recording their EODs with distributed electrodes (Fig. 1 B) — as has been suggested by Hagedorn and Heiligenberg (1985) and tested in the lab (Jun et al., 2013; Matias et al., 2015) and in the field (Madhav et al., 2018) — it is possible to track individual fish in their natural habitats without the need to tag them (Henninger et al., 2018). Such recordings pick up the EODs of all electric fish species present in the recording area. The frequency of electric organ discharges in wave-type fish is species- and individual-specific and, in constant environments, remains remarkably stable over many hours and days (Bullock, 1970; Moortgat et al., 1998), thereby providing individual frequency tags ideally suited for tracking individual fish.

**Figure 1:**
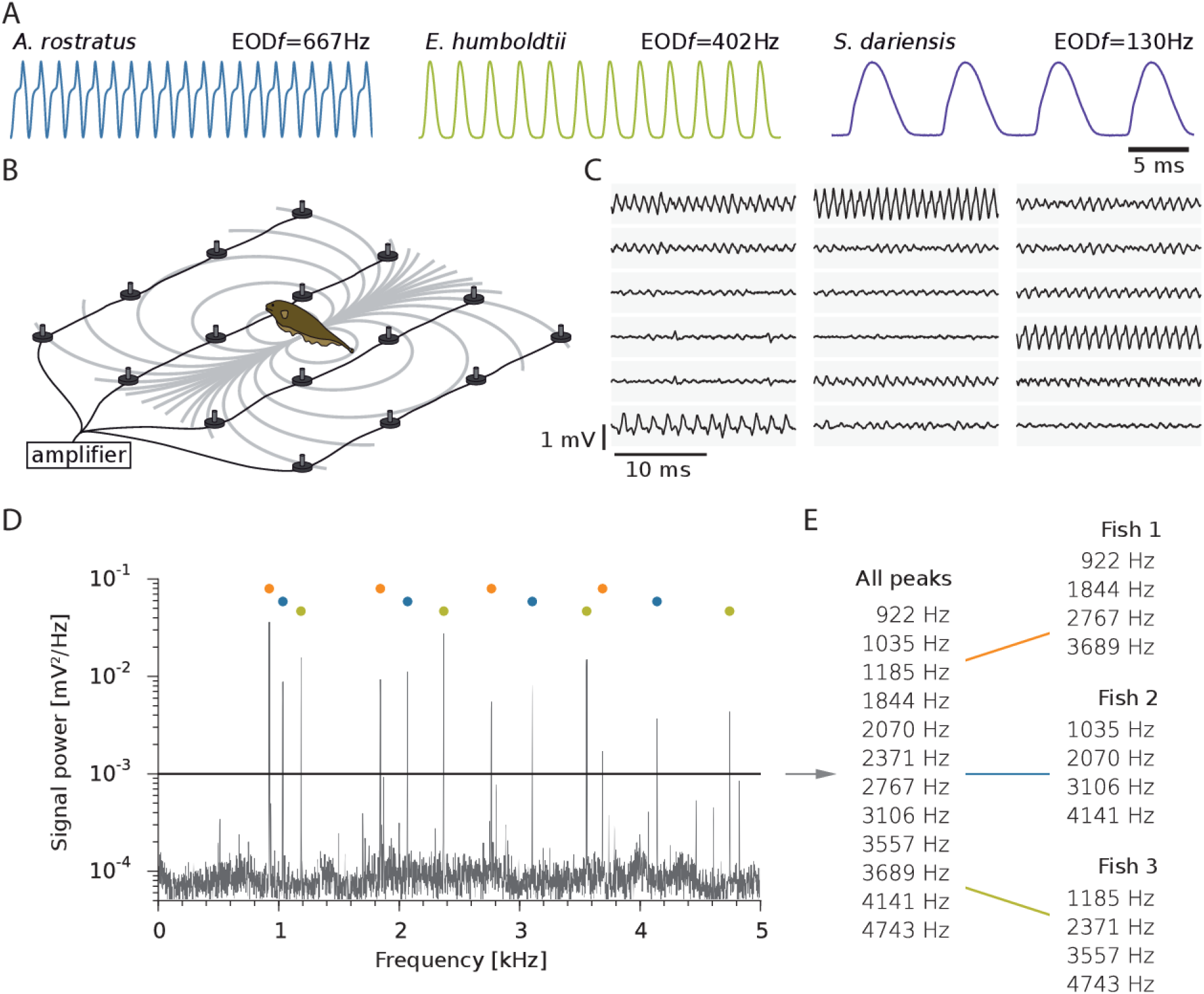
Recording untagged weakly electric fish with an electrode array. A) Example EOD waveforms of the three species of wave-type weakly electric fish present at our recording site. B) The electric organ discharge (EOD) of weakly electric fish generates a dipolar electric field (gray isopotential lines) that we recorded with an electrode array. C) An example of raw electrode data recorded in a stream in Darién, Panamá (Henninger et al., 2018). Each of the gray boxes shows the recorded voltage of one of the electrodes as they were arranged in the grid. In this example, three *Apteronotus rostratus* were concurrently present, a low-frequency female (lower left) and two high-frequency males (upper left and center right). The EOD of each fish was captured simultaneously by multiple electrodes. D) Power spectrum of the EODs of three concurrently present electric fish recorded on a single electrode. E) Peak detection generates a list of prominent frequencies (left), which are sorted into groups (color coded) of fundamental frequencies with their harmonics (right).

The EOD is an integral part of an active electrosensory system that the fish use for detecting prey (Nelson and MacIver, 1999), navigation (Fotowat et al., 2013), and communication (Smith, 2013). Previous field studies on the behaviors of these fish have been based on brief and focused EOD recordings with single electrodes (e.g., Lissmann and Schwassmann, 1965; Steinbach, 1970; Hagedorn, 1988; Friedman and Hopkins, 1996). Such studies revealed the presence, distribution, and sometimes movements of electric fish in a given habitat, but less detailed information about their interactions. Similarly, studies on single species of non-electric fish equipped with radio or ultrasonic transmitters are often restricted to few samples per day (e.g., Crook, 2004; Dawson and Koster, 2018). In contrast, overnight recordings of weakly electric fish at sub-second resolution with an electrode array (Fig. 1 B & C) revealed unexpected insights into the courtship behavior and its electrosensory implications (Henninger et al., 2018).

Here, we introduce the algorithms for identification and position estimation used by our automated approach for EOD-based tracking in the field (Henninger et al., 2018). We quantified the performance of three algorithms for position estimation using data obtained in a laboratory setting and from simulations. Using an array of 54 electrodes submerged in a small neotropical stream (240 × 150 cm at 30 cm spacing) we continuously recorded the electric activity of all electric fish passing through the array. We quantified EOD characteristics, activity and movement patterns of three species of wave-type gymnotiform fish (*Apteronotus rostratus*, *Eigenmannia humboldtii*, *Sternopygus dariensis*, Fig. 1 A) that were simultaneously present in our recordings.

## Materials and Methods

### Far-field measurement

We recorded the spatial amplitude distribution of the EOD’s electric far field potential in a large outdoor tank (3.5 × 7.5 × 1.5 m, *w × l × h*) at the Biocenter of the LMU, Martinsried, Germany. We used a 4 × 4 electrode array mounted on a PVC frame and spaced at 36 cm (108 × 108 cm, Georg Fischer GmbH, Albershausen, Germany). Electrodes were directly attached to monopolar headstages (1× gain). Signals of each electrode where amplified (100× gain), filtered (1st order high-pass filter 100 Hz, low-pass 10 kHz, npi electronics GmbH, Tamm, Germany), and digitized with 20 kHz per channel at 16-bit resolution. Water depth was adjusted to 60 cm, conductivity to 150 µS, and temperature to 23.5 °C. The electrode array was positioned at 30 cm water depth. A weakly electric fish (*A. albifrons*, 18 cm) was placed within the center of the electrode array on level with the electrodes, using a fish holder made of fine nylon mesh. During the measurement the fish did not change posture. In order to increase the spatial resolution we kept the position of the electrodes constant and instead adjusted the fish’s location. In this way we sampled a grid with a total of 108 locations in 18 *×* 6 steps resulting in a resolution of 2 *×* 4.5 cm (Fig. 2).

**Figure 2:**
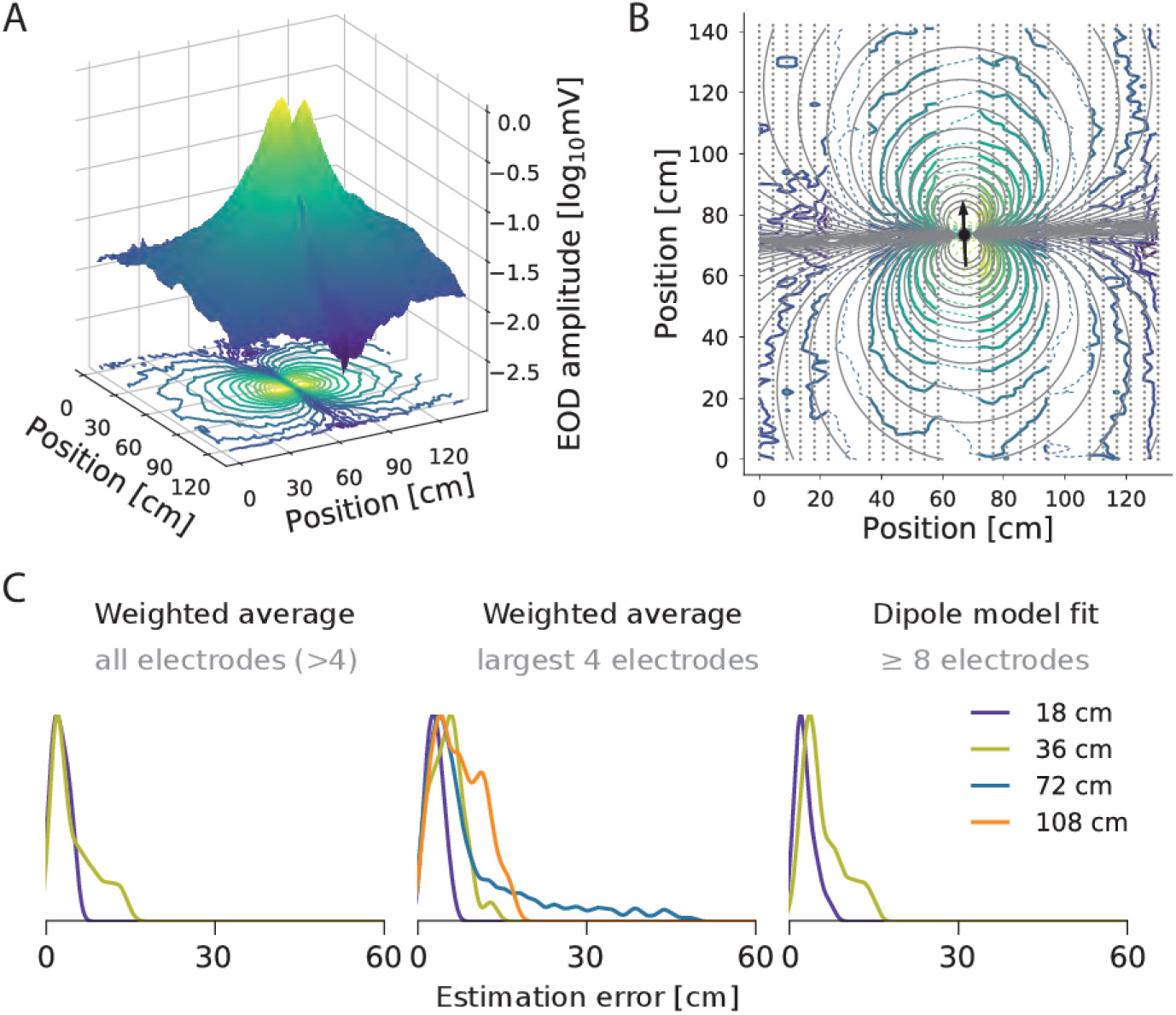
Spatial amplitude distribution of the EOD’s electric far field potential and localization errors. A) Absolute values of the EOD amplitude of an *A. albifrons* measured in a large tank in the horizontal plane. B) Dipole model (gray lines) fitted to the data shown in A (continuous contour lines). Interpolated data are indicated by dashed contour lines. The fish is located at the center and exactly in the plane of the electrode array and facing (arrow). C) The distribution of localization errors for different electrode spacings and algorithms as indicated applied to the data.

### Field site

Data of natural electric fish populations were recorded in Quebrada La Hoya, a narrow and slow-flowing creek, 2 km from the Emberá community of Peña Bijagual in the Tuira River basin, Province of Darién, Republic of Panamá (8°15′13.50″N, 77°42′49.40″W) in May 2012. At the recording site the water level ranged from 70 cm at the undercut bank to less than 20 cm at the slip-off slope. Water temperature and light intensity was recorded using a battery-powered data logger mounted close to the field site (HOBO Pendant Temperature/Light Data Logger, Onset Computer Corporation, USA). The water temperature varied between 25 and 27 °C on a daily basis and water conductivity was stable at 150 – 160 µS/cm. See Henninger et al. (2018) for more details.

### Field monitoring system

For the field recordings a custom-built, a 64-channel electrode and amplifier system running on 12 V car batteries (npi electronics GmbH, Tamm, Germany) was used (Fig. 1). Signals from stainless-steel electrodes were directly fed into low-noise headstages encased in epoxy resin (1× gain, 10×5×5 mm, Fig. 1 B) that measured the potential of the electrodes against the one of a common ground that was buried about 10 m downstream into the bank of the stream. After amplification and filtering by the main amplifier (100× gain, 1st order high-pass filter 100 Hz, low-pass 10 kHz) the signals were digitized with 20 kHz per channel at 16-bit using a custom-built low-power-consumption computer with two data-acquisition cards (PCI-6259, National Instruments, Austin, Texas, USA). Recordings were controlled with custom C++ software (https://github.com/bendalab/fishgrid) based on the comedi library for data acquisition on Linux (`comedi.org`).

We used 54 electrodes, mounted on a rigid frame (thermoplast 4 × 4 cm profiles, 60 % polyamid, 40% fiberglass; Technoform Kunststoffprofile GmbH, Lohfelden, Germany), and arranged in a 9 × 6 array covering an area of 240 × 150 cm (30 cm spacing, about two to three times the body length of the fish). The electrode array was submerged into the stream about 30 cm below the water surface and 40 cm above the sandy stream bed at the cut bank to less than 20 cm at the slip-off slope. The left column of electrodes was positioned below the washed out root masses of the cut bank (Fig. 4 A).

**Figure 3:**
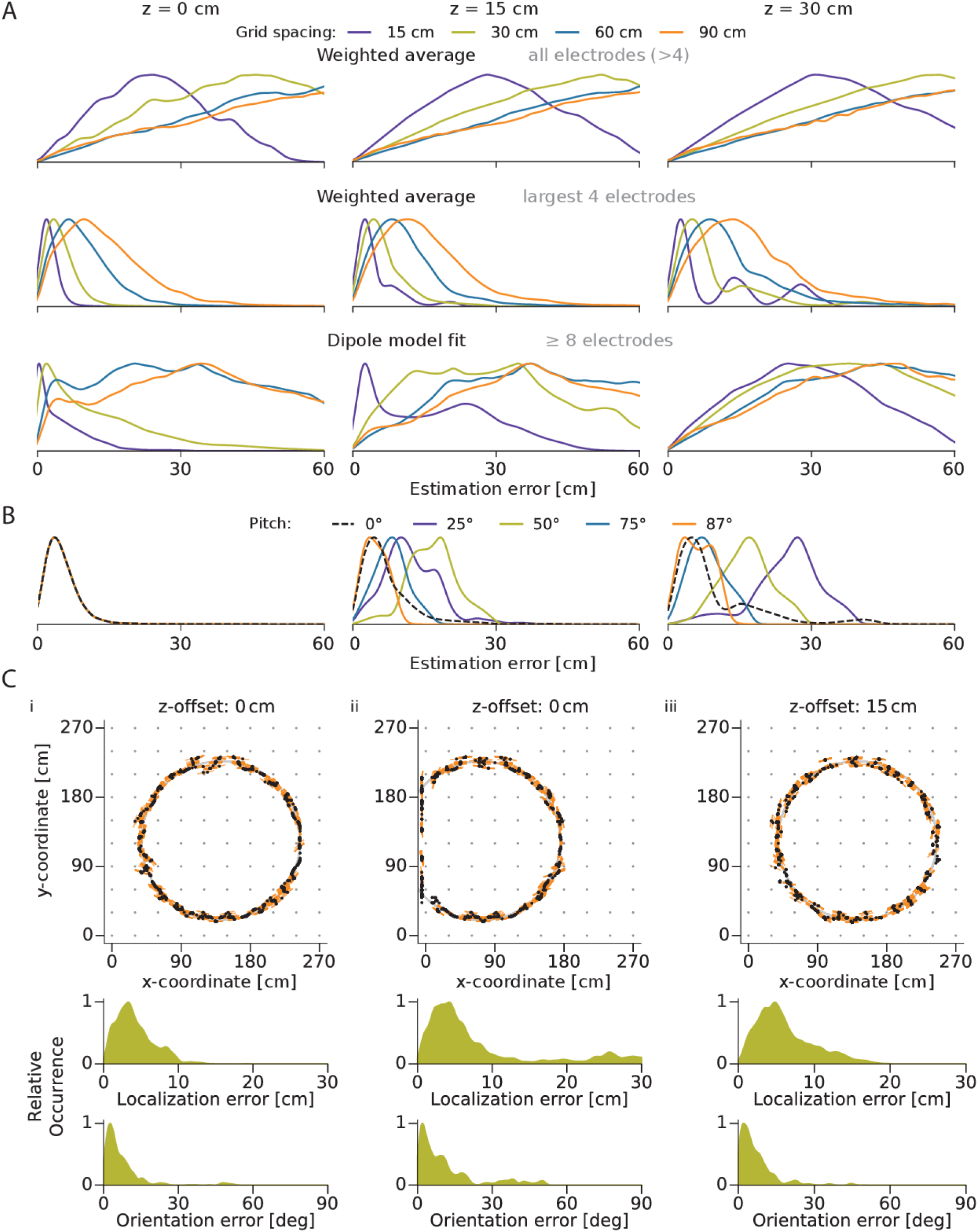
Performance of three algorithms for position estimation evaluated on simulated data. A) Distribution of localization errors computed for three different levels of the true static fish location above the plane of the electrode array (columns; 9 × 6 electrode layout). Only the weighted average taken over the four electrodes with the largest power consistently resulted in the smallest localization errors that stayed well below the electrode spacing. B & C) Further evaluation of the algorithm taking a weighted average over the four electrodes with the largest power for an electrode spacing of 30 cm. B) Distribution of localization errors for the same three *z* levels as in A computed from simulations of various pitch angles of the fish’s longitudinal body axis relative to the grid plane. C) Performance of position and orientation estimates of simulated moving fish. Top: The estimated, non-smoothed 2D-locations (black dots) and orientations (orange lines) were compared to the simulated fish locations and orientations (gray circle). Center and bottom: Distributions of errors in the estimation of localization and orientation. Ci) Simulated movement on level with the electrode array. Cii) Movement on level with the electrode array, but partially outside of the array. Errors include the difference to the known fish location and orientation for the time period the fish moves outside of the array. Ciii) Simulated movement 15 cm above the electrode array.

**Figure 4:**
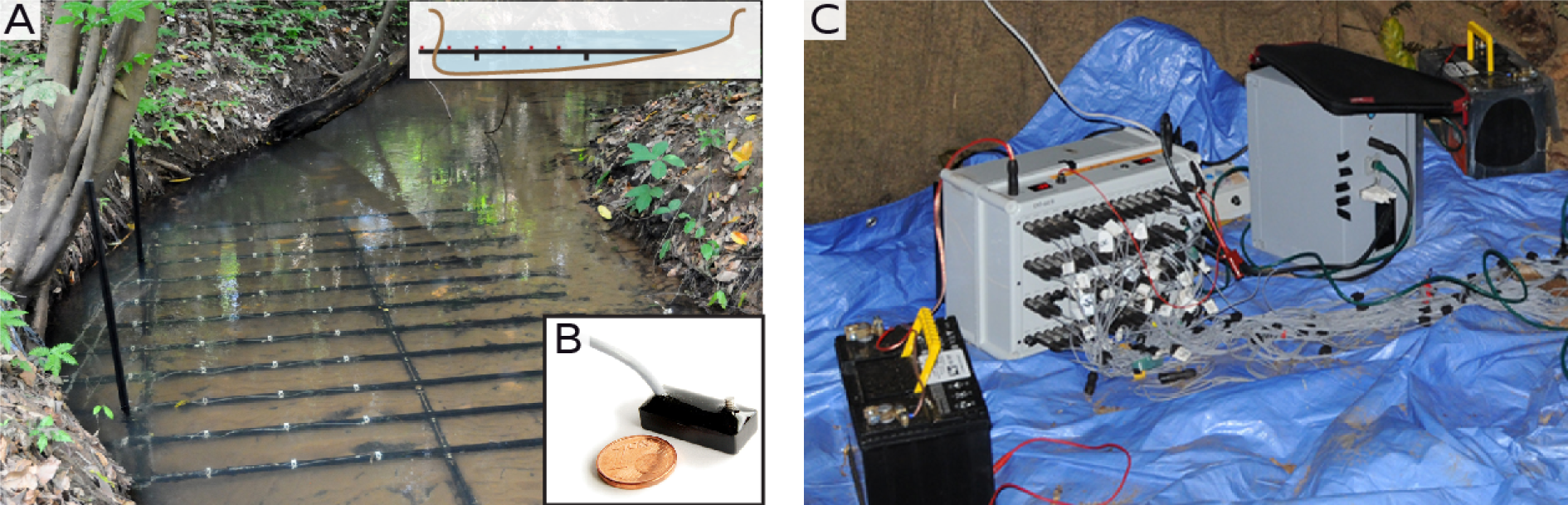
Recording site in a Neotropical stream and electrode array for recording electric fish. A) The 9 × 6 electrode array was submerged in a small stream in Darién, Panamá. White plastic holders keep the headstages (B) in place. Electrodes were positioned partly beneath the undercut banks, allowing to record electric fish hiding and interacting deep in the root masses. The schematic shows the stream’s approximate depths profile at the center of the array. Maximum water depth was 70 cm. B) Electrode headstage. The actual electrode is the stainless-steel screw. C) 64-channel amplifier (gray box to the left) and recording computer (right box) powered by two car batteries (black with yellow handle).

**Figure 5:**
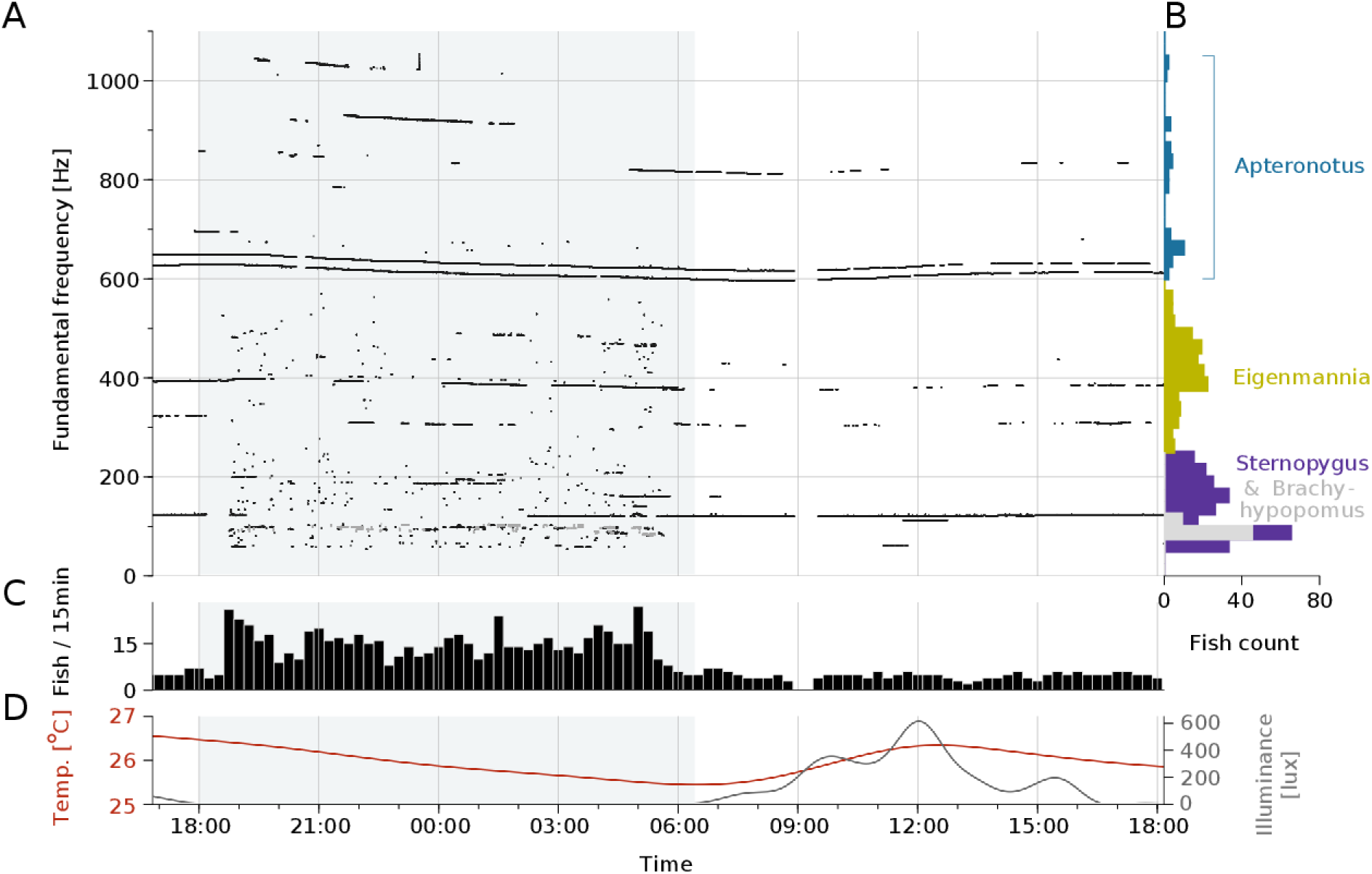
Diurnal activity pattern of a multispecies community of gymnotiform weakly electric fish. A) Fundamental frequency traces extracted from a continuous 25 hour recording (2012-05-10 to 2012-05-11). Each dot and horizontal line indicates the fundamental frequency trace of an individual detected electric fish. B) Histograms of the EOD*f*s shown in B separated into species. C) Temporal histogram of fish detections. Bin size is 15 minutes. D) Water temperature (red) and light levels (gray) during the recording period. Gray rectangles indicate nighttime. The interruption of the recording at 9am was caused by a switch of batteries needed to power the setup.

### Data analysis

Data were analyzed in Python 2.7 (www.python.org, https://www.scipy.org/, Hunter, 2007). Scripts and raw data (Panamá field data: 2.0 TB, same data set as in Henninger et al., 2018) are available on request. Data of the extracted EOD frequencies and position estimates are available at https://doi.org/10.12751/g-node.87584d. The core algorithms are accessible at Github under the GNU general public license (https://github.com/bendalab/thunderfish).

#### Individual identification

Individual fish were identified based on their EOD frequency and their EOD’s harmonic structure. The prime assumption for the EOD detection is that the EODs of wave-type electric fish possess a harmonic structure with at least three prominent components. For each electrode we calculated power spectral densities (PSD, 8192 FFT data points, 5 overlapping sub-windows, overlap 50%, total width= 1.22 s) in subsequent analysis windows (85 % overlap). In the log-transformed PSDs we detected peaks using a relative threshold (set to 1 dB) for the peak height (Todd and Andrews, 1999). The frequency resolution of the PSDs and peak positions was ∆*f* = 2.44 Hz.

For finding conclusive harmonic structures we started with the frequency *f*_*max*_ of the peak with the highest amplitude and checked whether harmonics at integer multiples of this frequency were present. As the fundamental frequency, *f*_1_, is not always the strongest frequency in an EOD’s PSD or might be even missing because of high-pass filtering, we checked for harmonic structures with fundamental frequencies at integer fractions of *f*_*max*_, i.e. *f*_1_ = *f*_*max*_/*n* for a small range of integers *n*≤4. Because of the discrete frequency resolution of the PSD, the fundamental frequency has an uncertainty of *f*_*tol*_ = ±∆*f*/2. In practice we set *f*_*tol*_ slightly higher (*f*_*tol*_ = ±0.7∆*f*) because peaks are distorted when riding on the flank of larger peaks. When checking for harmonics at frequencies *f*_*i*_ = *i* · *f*_1_ with *i* > 1, the corresponding frequency tolerances ±*i* · *f*_*tol*_ grow with the order *i* of the harmonics. Thus, the frequency tolerances get rather unspecific for higher harmonics, but at the same time subsequent harmonics of order *i* and *j* should also be separated by (*j* − *i*)*f*_1_ within a tolerance of 2|*j* − *i*|*f*_*tol*_. Having identified a potential harmonic of the fundamental frequency we used its frequency *f*_*i*_ to improve the estimate of the fundamental frequency via *f*_1_ = *f*_*i*_/*i*. Thus, by means of the harmonics, fundamental frequencies can be estimated with higher accuracy than the frequency resolution of the PSD. This updating of *f*_1_ was stopped as soon as a predicted harmonic was not present in the PSD.

The resulting group of harmonics was rejected if it contained less than three harmonics, if more than one of its frequencies was already contained in a harmonic group of another fundamental frequency, if more than a quarter of its harmonics were not detected in the PSD, or if no peak in the group was larger than an absolute threshold set to 50 µV^2^/Hz. Then, the group of harmonics was compared to the so far best group found for a given *f*_*max*_ and preferred if the sum of its peak amplitudes was larger and the number of missing harmonics was lower. This procedure was repeated until all peaks in the power spectrum of a certain amplitude that was higher than the detection threshold were considered. As a final step, harmonic structures with fundamental frequencies outside the expected EOD *f* spectrum (40 Hz –– 1500 Hz), or at the mains hum at 60 Hz were discarded.

The algorithm for retrieving fundamental frequencies fails, if peaks in the power spectrum are smeared out because of rapid changes in EOD frequency caused, for example, by gradual frequency rises (Turner et al., 2007), or by electric noise pulses (Hopkins, 1973). Also, if peaks approach the noise floor they will be missed and the corresponding harmonic group cannot be retrieved any more. Both issues did not pose a problem in our data set. The distinct harmonic structures of EODs of different species (e.g. Fig. 6) did also not affect detection performance.

**Figure 6:**
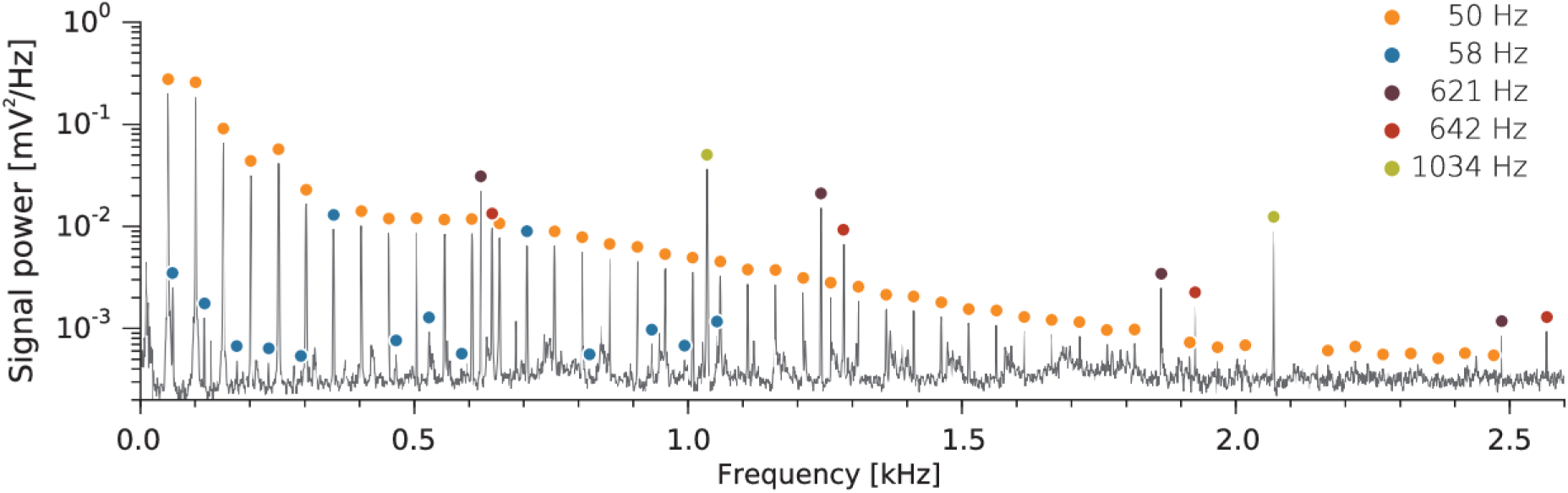
Example power spectrum of the averaged signal of the array recorded in Darién, Panamá (at 20:44:18 local time in Fig. 5). The EODs of five concurrently present electric fish of two species clutter the power spectrum with many intermingled peaks. Two low-frequency *Sternopygus dariensis* marked with orange and blue dots (< 200 Hz) and three high-frequency *Apteronotus rostratus* (> 580 Hz, purple, red, and green dots) can be extracted from the power spectrum. Detected EODs are displayed on the right. Colored markers indicate the EODs’ harmonic structures.

#### Temporal tracking of electric fish

In a second step, detected EOD frequencies were connected to previous data segments in order to track individual fish over time. If a fish with an EOD *f* differing by less than 10 Hz was found in the previous detections, the new data was added, otherwise the fish was treated as a new candidate. If this candidate was detected robustly in the following analysis windows, typically over several seconds, it was marked as a confirmed fish detection. Otherwise, it was discarded as a false detection. Confirmed fish detections remained in an ‘active’ state for a short time period (here: 10 minutes) after which they were set to ‘passive’ state. Later detections of a similar EOD frequency were treated as new detections.

A critical problem for temporal tracking wave-type electric fish are EOD frequencies that approach, or even cross, each other. This is not a particular problem in the data set introduced here, but could easily occur at larger fish densities or with social fish like *Eigenmannia* (Henninger, 2015; Madhav et al., 2018). This issue can, to a certain extent, be be resolved by taking the spatial distribution of power at a given frequency into account (Madhav et al., 2018).

#### Estimation of location and orientation

For each detected fish the electrode’s voltage traces were bandpass-filtered (Butterworth filter, 3rd order, 5× multi-pass, ±7 Hz width) at the fish’s EOD*f*. For each passband the signal amplitude was estimated each 40 ms using a root-mean-square filter over 10 EOD cycles multiplied by 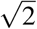. These amplitudes were then used to estimate the location and orientation of each fish. In the following we describe the algorithms tested for fish localization. We first introduce two variants of a weighted spatial average, before we discuss an algorithm based on a dipole-model. Note that all three algorithms only estimate the orientation of the body axis, they do not discriminate between head and tail.

##### Weighted spatial averages

Two variants of estimating 2D fish location and orientation were based on weighted spatial averages. The fish position 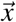 was estimated from *n* electrodes *i* with the largest envelope amplitudes *A*_*i*_ at position 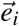 as a weighted spatial average, given by

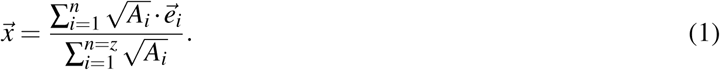

For the variant that used all electrodes, *n* was set to the number of electrodes of the electrode array. For the other variant that used only the four electrodes with the largest EOD amplitude, the position was computed only, if at least 2 electrodes with amplitudes greater than 15 µV were available. If at least 4 electrodes with amplitudes greater 1 µV were available, *n* = 4, otherwise *n* = 2.

The EOD of a fish can cause a very large amplitude on nearby electrodes because of the electric field’s reciprocal dependence on distance. This effect results in a relatively large localization error, if a simple weighted spatial average is used, because the position estimate is pulled towards the strongest electrode. The localization error is reduced by using the square-root of the EOD amplitude, 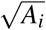, as a weight, which reduces the impact of electrodes with large EOD amplitudes.

For approximating fish orientation, we first divided the electrodes into two subgroups of opposite polarity. Because the EOD amplitudes are extracted as absolute values, polarity information of the EOD on the respective electrodes is missing and has to be estimated in an additional analysis step. The polarity of the electrodes was determined by calculating the correlations of the electrodes’ bandpass-filtered voltage traces (40 ms windows) relative to that of the electrode with the largest amplitude. Electrodes with correlations larger than +0.9 were assigned to one group and correlations smaller than −0.9 to the other. If both groups contained at least 4 electrodes, each group’s center was estimated by calculating the weighted spatial average, Eq. (1). The direction of the vector connecting the centers of the two groups was an estimate of the orientation of the body axis of the fish.

##### Dipole model

The potential generated by an ideal electric dipole at a distance *r* and an angle ϕ measured against the dipole moment is given by

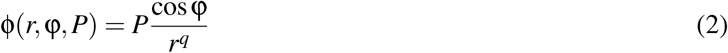

where the exponent *q* = 2 and *P* is an amplitude factor that absorbs the dipole moment (directly proportional) and the permittivity of the medium, i.e. the water conductivity (inversely proportional). The electric organs of weakly electric fish can be approximated as constant current sources (Knudsen, 1975) and therefore *P* is proportional to the current produced by the electric organ and inversely proportional to the conductivity of the water (Madhav et al., 2018).

For estimating the position, orientation, and EOD amplitude of a fish we fitted the absolute value of the dipole model Eq. (2) to the EOD amplitudes recorded by the electrode array. This is a numerically difficult minimization problem, because of the large number of local minima between the singularities at the positions of the electrodes. We therefore introduced a regularizer α in the denominator, to remove these singularities and make the problem numerically more stable. We also treated the exponent of the power law as a free parameter *q* to phenomenologically account for the compression of the electric field by the non-conducting water surface and bottom (Knudsen, 1975). Together, these modifications yield the regularized dipole function, given by

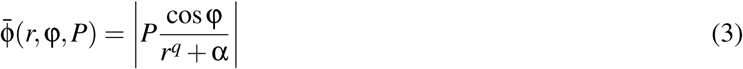

A weighted spatial average over all available electrodes was used to compute initial values for the optimization. The optimization was performed over two iterations using stepwise decreasing values for α (10^−2^ and 10^−3^), and using the result of the first iteration as initial values for the last. The successful optimization directly yielded the dipole’s location, orientation, its amplitude factor, and the potential distribution’s effective power law. Since we were optimizing a function with 8 parameters, at least 8 data points, i.e. amplitude measurements from at least 8 electrodes, were necessary for a successful optimization.

##### Simulations

To study the performance of the three proposed localization methods under ideal conditions, we simulated large electrode arrays (40 × 40) with spacings of 15 – 90 cm and sampled many fish locations and orientations at the array’s center quadrant (20 × 20 steps xy-resolution; 3 elevations; orientations between 0 – 180° with 4.5° resolution; 16400 samples/spacing, total). This configuration allowed for neglecting effects caused by the limits of the electrode array. Data were simulated with a dipole model using EOD field parameters extracted for *A. albifrons* and a power law with exponent −1.64 (Fig. 2 B).

In order to estimate the detection performance of our analysis chain for moving fish, we simulated the EODs of moving fish (*v* = 10 cm/s) generated with a horizontally oriented dipole model, Eq. (3) and electric field parameters extracted from our EOD field measurements. Electrodes were configured in a large array with 10 × 10 electrodes with a spacing of 30 cm, i.e. the same spacing as used at our field site. Data were generated using the same sample rate of 20 kHz as for the field data. For each sample we calculated the EOD’s potential on each electrode and multiplied it with the current value of the EOD. The simulated fish moved a full circumference of a circular trajectory (103.3 cm radius), always with the circle positioned slightly off-center in relation to the electrode array’s grid in order to sample many different fish-to-electrode configurations. The noise floor in the field recordings was at about 1 µV. We therefore excluded electrodes with amplitudes below 1 µV from the position estimate to get comparable performances to the field situation.

#### Movement patterns

Searching for directed movements through the recording area, we split all recorded fish movement traces at detection gaps of 20 s and more. As directed movements we classified trajectories that entered the electrode array in one third and left the array at the opposite third in the direction of the stream flow (the *y*-coordinate of the electrode array). We did not impose any time limit. Average upstream (*v*_*up*_) and downstream (*v*_*down*_) swim speeds were computed as the slope of a straight line fitted to each *y*(*t*) trace in the central half of the electrode array (60 cm < *y* < 180 cm). Using a larger or smaller part (±15 *cm*) of the electrode array did not change the results.

The measured upstream speed of a fish, *v*_*up*_ = *v*_*fish*_ − *v*_*stream*_, is the swimming speed of the fish, *v*_*fish*_, reduced by the water velocity of the stream, *v*_*stream*_. Likewise, the measured downstream speed is *v*_*down*_ = *v*_*fish*_ + *v*_*stream*_. By subtraction we get an estimate of the water velocity

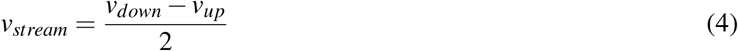

and by addition an estimate of the swimming speed of the fish

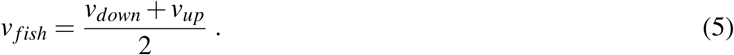

Since we did not know exactly which downstream swimming fish corresponded to which upstream swimming fish (the EOD frequency changes with temperature), we used the median speeds for the calculations.

Assuming the fish are constantly swimming with *v*_*fish*_ relative to the water body we would like to estimate how far a fish was able to swim upstream during one night if it returns after the time ∆*t*_*total*_. The fish can spend the time ∆*t*_*up*_ for swimming upstream with *v*_*up*_ and the time ∆*t*_*down*_ with *v*_*down*_ for swimming downstream. To be back in time, ∆*t*_*total*_ = ∆*t*_*up*_ + ∆*t*_*down*_ and the covered distances *x* need to match: *x* = ∆*t*_*up*_*v*_*up*_ = ∆*t*_*down*_*v*_*down*_. From these two conditions follows for the distance covered by the fish

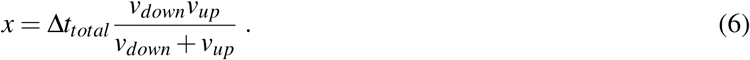

For the time ∆*t*_*total*_ we used the difference between the median times of upstream and downstream movements.

As the distance from the undercut bank we simply used the *x*-coordinates of the movement traces from the same segments as used for estimating swim speed.

#### EOD field characteristics

The far field of the electric field generated by weakly electric fish approximates that of an ideal dipole Eq. (2). The absolute potential φ at a given distance *r* varies because of its angular dependence on ϕ via the cosine term. Taking only the maximum measured values for each distance cancels out the angular dependence and we are left with a power law

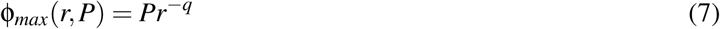

to be fitted to the data. This allowed for a robust estimate of the amplitude factor and the exponent of the power law decrease of electric field amplitude. Note that in shallow water the exponent can be smaller than two (Fotowat et al., 2013).

To extract the maximum amplitudes, amplitudes obtained from bandpass-filtered data were binned logarithmically over distance and for each bin a fixed fraction of largest amplitudes (here 5%) were extracted and averaged. Eq. (7) was then fitted to the obtained amplitudes. Errors introduced by the inaccuracy of location estimation were particularly noticeable at small distances. At large distances the data hit the noise floor (Fig. 10 A). We therefore excluded distances below 20 cm and above 100 cm from the fit.

For an estimation of the detection ranges we first computed the electric field strength *E*(*r*) as the spatial derivative of Eq. (2) to

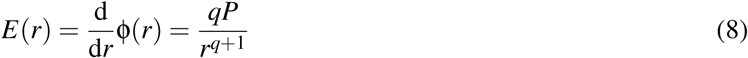

where we have set cosϕ = 1. Given the sensitivity *E* we get for the maximum detection range

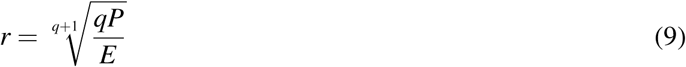

#### Transects

We recorded positions of wave-type weakly electric fish along a 160 m long transect directly upstream of the electrode array on 2012-05-11 at around noon and on 2012-05-13 in the afternoon. We walked along the transect and located the fish by means of a fish-finder, a 30 cm long custom-made dipole electrode connected to an audio amplifier (Mini Amplifier-Speaker, RadioShack). For each fish we recorded the EOD for about 20 s (24 kHz, XL2 Audio and Acoustic Analyzer, NTi Audio GmbH, Essen) and its position along the stream. From the recordings we extracted EOD frequencies using our thunderfish software (https://github.com/bendalab/thunderfish). Sometimes the same fish was present on several recordings, in that case we only counted the fish in the recording where it had the largest power and removed it from the other ones. We then counted the number of fish present in the transect for each species separately based on their EOD frequency. Division by the transect length resulted in average fish densities. The smooth EOD waveforms shown in Fig. 1 A have been generated by averaging field-recorded EODs of the respective species 100-times each. This removed smaller EODs from concurrently present pulse- and wave-type fish.

## Results

### Identification of individual electric fish by EOD frequency

Individual wave-type fish differ in the frequency of their EODs. Extracting the EOD frequency (EOD*f*) from the raw data thus allows identification of individual fish. This conceptionally simple task is complicated by the fact that often more than a single fish is picked up by an electrode (Fig. 1 C). Furthermore, because EOD waveforms of wave-type fish are distorted sine waves, the resulting frequency spectra of individual fish contain harmonics, i.e., peaks at multiples of the fundamental EOD*f*. Together this results in complex frequency spectra where peaks originating from individual fish are intermingled (Fig. 1 D).

We solved this problem by analyzing the frequency spectra for periodically occurring peaks (i.e., harmonics) and assigning them to individual fish (Fig. 1 D, see methods). In a next step, individuals detected on single electrodes were matched with those of similar fundamental frequencies found on other electrodes, and finally matched over sequential time steps to generate temporal consistency. This algorithm allowed us to robustly and automatically analyze datasets from at least six different species (*Apteronotus albifrons, A. leptorhynchus, A. rostratus, Sternopygus dariensis, Eigenmannia humboltii and Sternarchorhynchus spec.*; Henninger, 2015; Henninger et al., 2018).

### Dipole-like far-field of EODs

Crucial for studying behavior is knowledge about each individual’s position and movements. For determining effective layouts of the electrode array and for choosing a suitable localization approach, we extended previous studies (Heiligenberg, 1973; Knudsen, 1975; Rasnow et al., 1993; Rasnow and Bower, 1996) and measured the electric field’s amplitude distribution of an *Apteronotus albifrons* over longer distances in a large rectangular outdoor tank. As expected, the measurements resembled the field created by an electric dipole (Fig. 2 A). The EOD amplitude was attenuated with distance from the fish and was modulated as a function of the angle relative to the animal’s body axis. A modified dipole, Eq. (3), fitted to the data resulted in a good description of the measured EOD amplitudes (Fig. 2 B) with exponent *q* = 1.63 and amplitude factor *P* = 29 mV cm^*q*^.

### EOD-based localization of electric fish

Next, we evaluated the performance of three algorithms for estimating fish position. Two simple estimates of 2D fish location and orientation were based on spatially averaging electrode positions weighted by the EOD amplitudes measured at the electrodes. In one version we used all available electrodes to compute the fish location, and in another version only the four electrodes with the largest EOD amplitudes. A third algorithm fitted a dipole model Eq. (3) to the measured EOD amplitudes.

When applied to the measurements from Fig. 2 A, B, where the fish is positioned in the center and at the vertical level of the electrode grid, all three estimators performed reasonably well with median position errors of about 5 cm for inter-electrode distances ≤ 36 cm (Fig. 2 C). Increasing the distance between electrodes to 108 cm resulted in a minor increase of the median estimation error to about 10 cm.

In reality, however, fish are not always located at the center of the electrode array. We therefore tested the performance of the three position estimators in simulations in which we varied the static position, orientation, and level above the electrode array as well as the electrode spacing using a realistic 9 × 6 electrode layout (Fig. 3 A). Fitting the dipole model required narrow electrode spacing and performance deteriorated dramatically for fish swimming outside the plane of the electrode array. Using data from all electrodes of the array for the weighted spatial-average estimate resulted in errors of the same size as the electrode spacing. Only when using the four electrodes with the largest EOD amplitudes was the position estimate of the weighted spatial average largely independent of the level above the electrode array and the error was much smaller than the electrode spacing — although this estimate does not relate to the underlying physics of the electric field. We therefore used this measure for further analysis.

Gymnotiform fish commonly tilt their body axis during feeding, explorative behaviors, and social interactions. If the fish is on level with the electrodes, various pitch angles of the body axis do not influence localization performance (Fig. 3 B). Yet, if the pitched fish is offset from the electrode array, the estimation error increases with the offset and in particular with small pitch angles. Note, that the estimation errors introduced by pitch stay below the ones of the other two algorithms (weighted average of all electrodes or dipole fit) for zero pitch (compare to Fig. 3 A).

### Position and orientation estimates of moving fish

We also studied how position estimation performs with moving fish. A fish moving with a speed of 10 cm/s (Nelson and MacIver, 1999) along a circular trajectory was simulated. Again, the median of the position errors was clearly below 10 cm (Fig. 3 Ci), which corresponds to the length of a small-sized, mature *Apteronotus rostratus* (Meyer et al., 1987). Likewise, the medians of the orientation errors were small and well below 15°. The weighted spatial average was unable to follow fish positions outside the boundaries of the electrode array. Therefore the error of the position estimate increased as soon as the fish trajectory extended beyond the electrode array (Fig. 3 Cii). For fish trajectories vertically offset from the grid plane, the median and spread of the localization error increased slightly, while the orientation error remained almost unchanged (Fig. 3 Ciii). In summary, the simulations demonstrated that the algorithm computing a spatial average of the four electrodes with the largest signal was suited for tracking electric fish moving within the electrode array’s limits with uncertainty below an adult fish’s body length.

### Diurnal activity patterns in the natural neotropical habitat

We applied our EOD tracking system in a small neotropical stream using an array of 54 electrodes to continuously record the electric activity of all electric fish passing over the array (Darién, Panamá; Fig. 4). We quantified EOD characteristics, activity and movement patterns of three species of wave-type gymnotiform fish (*Apteronotus rostratus, Eigenmannia humboldtii, Sternopygus dariensis* that were simultaneously present at our study site Fig. 4 A). The fundamental frequencies extracted from 25 hours of almost continuous recording demonstrate the richness and complexity of EOD frequencies present in a natural habitat of gymnotiform weakly electric fish (Fig. 5 A). In this example we registered 461 EOD detections, i.e. fundamental frequencies that were tracked continuously with possible interruptions of less than 10 minutes (see methods). This temporal tolerance entails that the number of detections was likely higher than the total number of individuals detected: when the same fish left and reentered the recording area over the course of the recording, separated by a gap of more than 10 minutes, it was treated as the detection of a new fish.

A histogram of the frequencies of the EOD detections revealed three distinct frequency ranges corresponding to three wave-type species (Fig. 5 B): *Apteronotus rostratus* occupied the highest frequencies from ~580 to 1100 Hz. Right below were *Eigenmannia humboldtii* between ~200 and 580 Hz. *Sternopygus dariensis* covered the lowest frequencies (~40 – 220 Hz) and shared its frequency range with the pulse-type fish *Brachyhypopomus occidentalis*. We confirmed all classifications by inspection of the EOD waveforms (Fig. 1 A).

The number of EOD detections per 15 minute time bin was significantly larger during the night than during the day (*p* = 1.1 × 10^−22^, Welch’s t-test; Fig. 5 C), directly demonstrating the nocturnal activity of weakly electric fishes (compare to illumination levels in Fig. 5 D). EOD frequency is known to be sensitive to water temperature (e.g., Coates et al., 1954; Dunlap et al., 2000). Therefore, diurnal changes in EOD frequency were expected given the temporal variation in water temperature (Fig. 5 D, e.g., the two fish right above 600 Hz in panel A).

### Multispecies EOD interactions

Power spectra of the recordings often contained signatures of several fish, often of different species (Figs. 5 A). The example shown in Fig. 6 shows three *Apteronotus* and two *Sternopygus* simultaneously present on a recording electrode. When looking at the reconstructed movements, we also found many scenes in which fish of different species were simultaneously present and in close proximity of each other (Fig. 7 A). The simultaneous presence of three wave-type species gave rise to complex interactions of their EOD waveforms.

**Figure 7:**
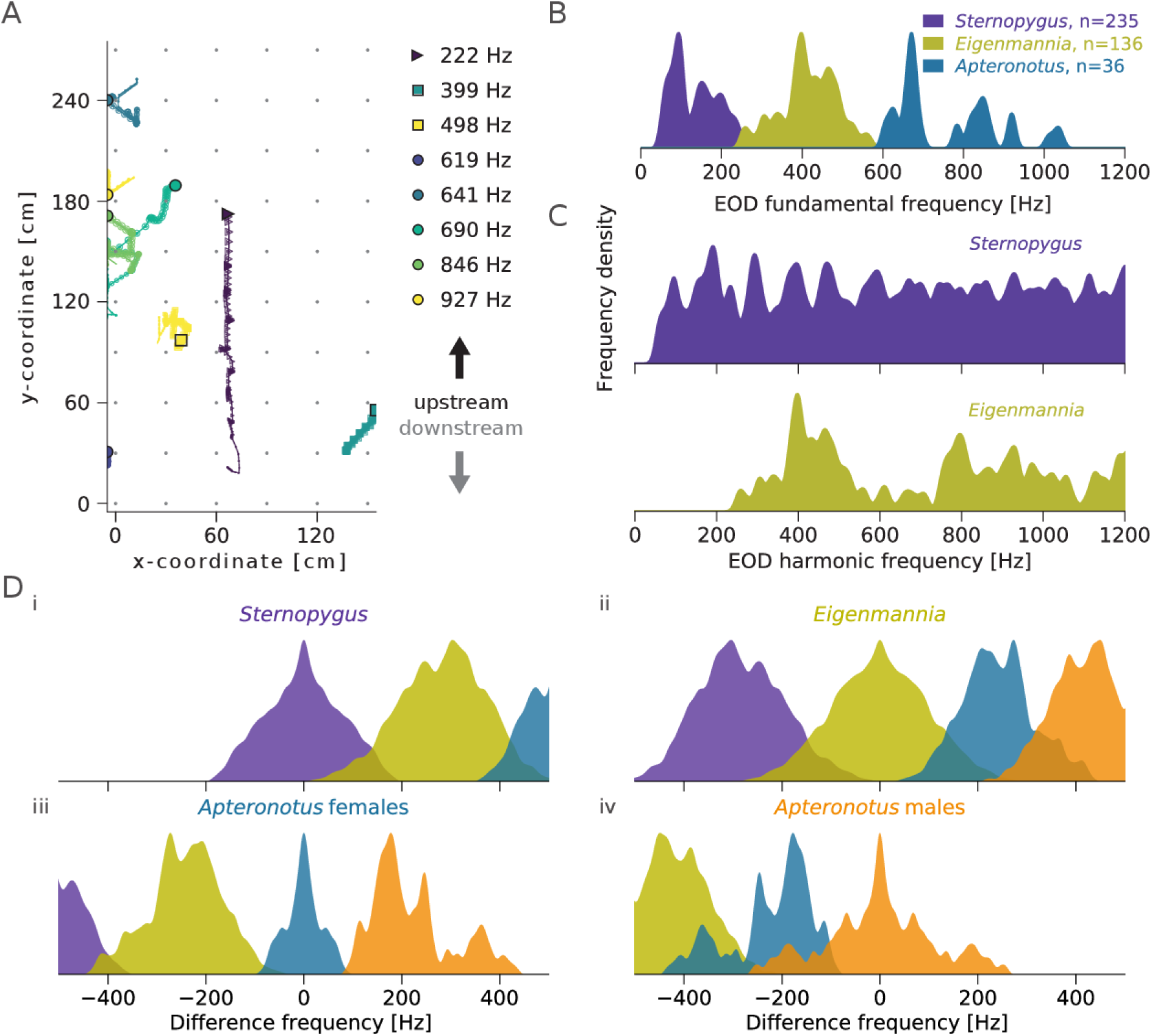
EOD interactions arising in a multispecies community of wave-type gymnotiformes. A) Example of concurrently present individuals of three sympatric species of electric fish recorded in the field (at 2012-05-10, 21:26:36). Legend indicates species and EOD fundamental frequency (triangle: *Sternopygus dariensis*, squares: *Eigenmannia humboltii*, circles: *Apteronotus rostratus*), tails indicate positions of the last 5 seconds. B) Distinct ranges of fundamental EOD frequencies for the three species recorded on the electrode array. Same as in Fig. 5 B, but histograms of each species are normalized to their respective maximum value. C) Distribution of fundamental EOD frequencies and their harmonics demonstrate that harmonics of lower-frequency species like *Sternopygus* and *Eigenmannia* extend well into the frequency ranges of higher-frequency fish (compare with panel B). D) Histograms of difference frequencies computed as the difference between all fundamental EOD frequencies of one species (colored histograms) and the fundamental EOD frequencies of another species (indicated by the title of each panel). Because of the considerable overlap between the histograms, it seems unlikely that the fish can differentiate between species based solely on EOD frequency differences.

First, harmonics of EODs with significant power easily extended into the frequency bands of species with higher fundamental frequencies. For example, the harmonics of a *Sternopygus* were, at times, of similar power as the fundamental frequencies of *Apteronotus* (Fig. 6). Whereas the fundamental frequencies of the three wave-type electric fish species were separated (Fig. 7 B), the harmonics of the lower-frequency fish *Sternopygus* and *Eigenmannia* clearly extended into the frequency range of the respective higher-frequency fish, i.e. *Eigenmannia* and *Apteronotus* (Fig. 7 C).

Second, the superposition of the EODs of two fish results in a beat, a periodic amplitude modulation, which is encoded by the electrosensory system (Bastian, 1981). The frequency of the beat is given by the difference between the fundamental EOD frequencies of the two fish. In contrast to the absolute fundamental EOD frequencies, these frequency differences are ambiguous, i.e. they cannot be used by an individual fish to unequivocally determine the species of another fish (Fig. 7 D). For example, a positive frequency difference perceived by a *Sternopygus* could have been induced by another *Sternopygus* or by a low-frequency *Eigenmannia* (Fig. 7 Di). Negative frequency differences perceived by an *Apteronotus* male could have been induced by either a rivaling male, a conspecific female, or even an *Eigenmannia* (Fig. 7 Div).

### Movement patterns

The ability to track individual movements enabled us to study species-specific movement patterns. Many of the detected fish stayed only for a short time within the electrode array (Fig. 5 A). In fact, they often simply swam straight through the array. From the 461 electric fish detections over 25 hours, a subgroup of 173 detections showed directed movement, i.e. a fish entered the electrode array on one end and left it at the opposite end in less than one minute (23 ± 10 s; range: 10 to 58 s), either with or against the direction of water flow (Fig. 8).

**Figure 8:**
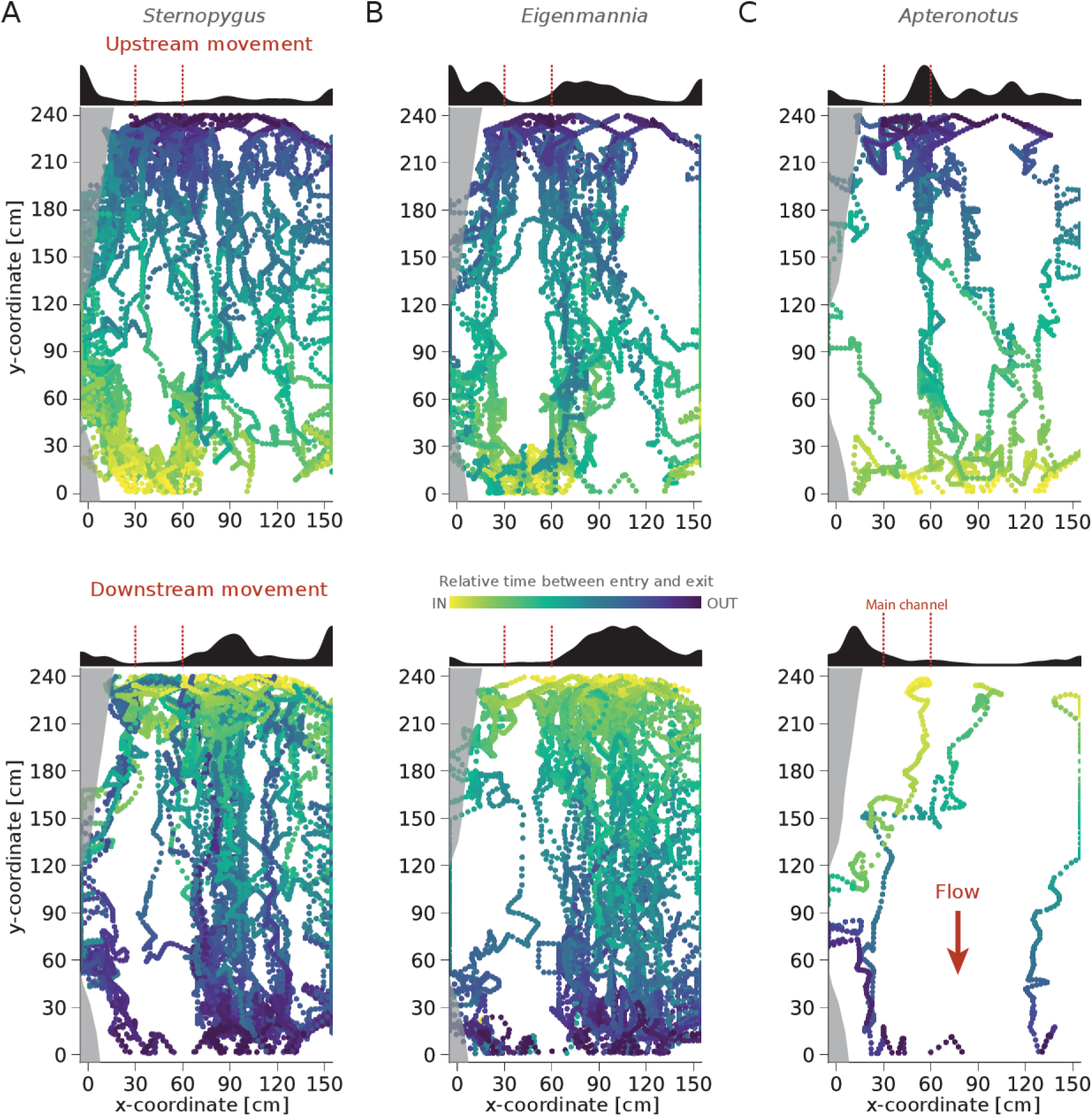
Trajectories of directed movements in the field. Top panels: upstream movements, bottom panels: downstream movements. Columns: A) *Sternopygus dariensis* (*n* = 98), B) *Eigenmannia humboldtii* (*n* = 65), C) *Apteronotus rostratus* (*n* = 10). Color encodes relative time between entering (yellow) and leaving (purple) the recording area. Shown are trajectories of all 173 directed movements traversing the recording area extracted from the 25 hour recording shown in Fig. 5. Movement traces have been smoothed with a running average (width=200 ms). Water flows from top to bottom (arrow in C). Histograms on top of the panels show distribution of fish locations along the *x*-coordinate of the traces computed for the center area (60 cm < *y* < 180 cm). The cut bank is indicated by gray shading, the main channel with the strongest water flow by red vertical lines on the histograms (compare to Fig. 4 A).

Movement activity set on sharply after nightfall. Individuals of all species had a strong tendency to move upstream in the first half of the night and downstream in the second half, a pattern most pronounced in the larger populations of *Sternopygus* and *Eigenmannia* (Fig. 9 A). Most fish were swimming solitarily through the grid. Only 8 % of 98 *Sternopygus* detections, 6 % of 65 *Eigenmannia* and none of 10 *Apteronotus* were swimming simultaneously with a conspecific into the same direction through the electrode array.

**Figure 9:**
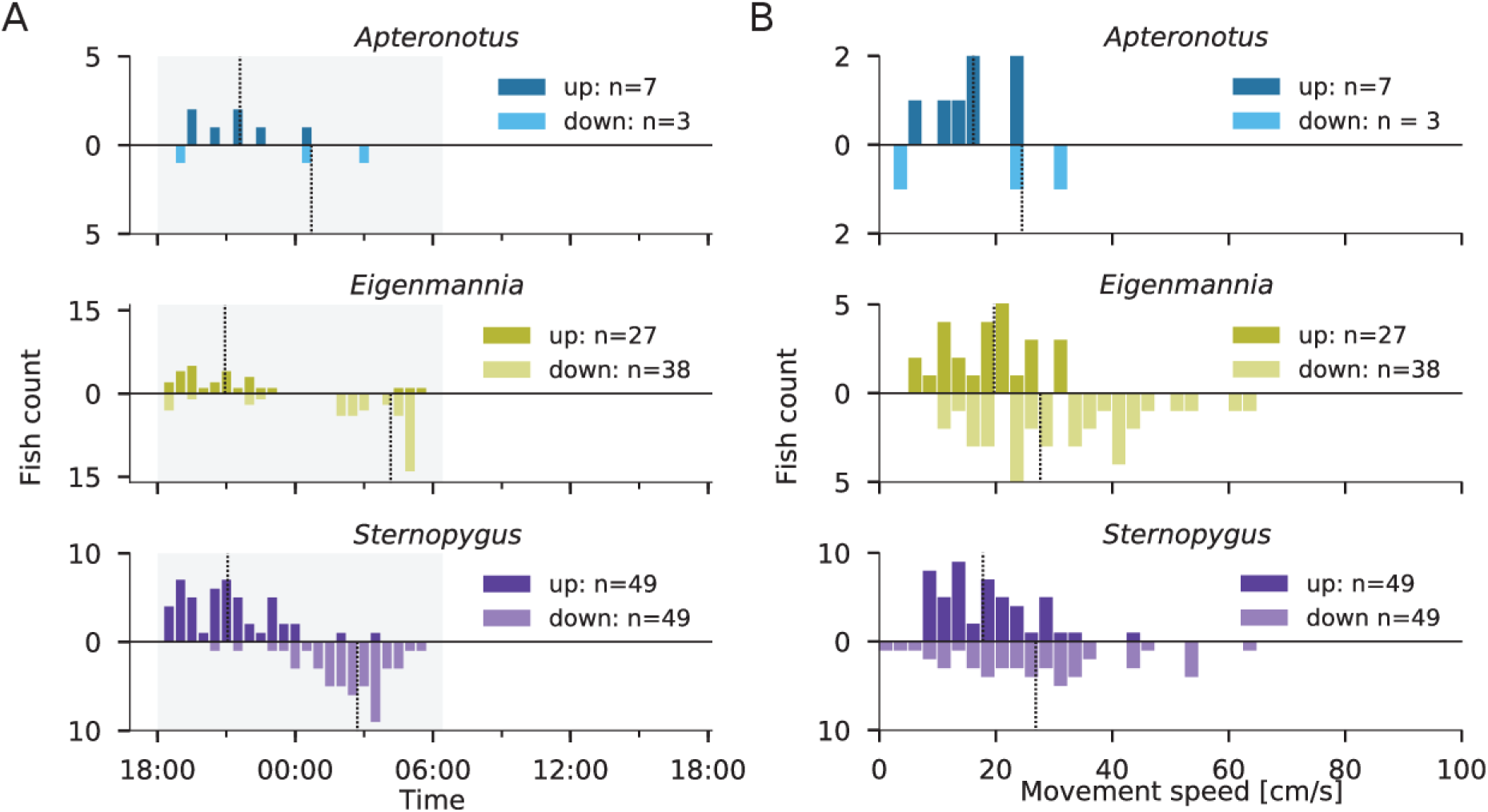
Statistics of fish traversing the electrode array. A) Time of occurrence of directed upstream and downstream movements of *Apteronotus, Eigenmannia*, and *Sternopygus*. The fish tended to swim upstream during the first half of the night and downstream in the second half. Nighttime is indicated by the gray background. B) Corresponding average movement speeds along the *y*-coordinate of the electrode array. Dashed lines indicate median times and speeds. Histograms are based on the trajectories shown in Fig. 8.

Upstream swimming *Sternopygus* and *Eigenmannia* showed a preference to swim along the root masses (on the left side of each panel of Fig. 8): 39 % of 49 *Sternopygus* (Poisson test: *p* < 0.01) and 26 % of 27 *Eigenmannia* (*p* = 0.18) moving upstream stayed within 30 cm of the undercut bank. In contrast, 92 % of 49 *Sternopygus* (*p* ≪ 0.001) and 95 % of 38 *Eigenmannia* (*p* ≪ 0.001) moving downstream stayed farther than 60 cm away from the undercut bank, preferring slower waters inside the meander bend. Neither up- nor downstream swimming fish were swimming in the main channel between 30 to 60 cm from the undercut bank (*p* < 0.01 for *Sternopygus* and *Eigenmannia*).

We quantified the average movement speeds along the stream (*y*-coordinate of the electrode array) of the tracked individuals (Fig. 9 B). *Sternopygus, Eigenmannia*, and *Apteronotus* were moving upstream with a median speed of *v*_*up*_ = 17.8, 19.7, 16.2 cm/s, respectively. Later in the night they were swimming downstream with *v*_*down*_ = 26.9, 27.7, 24.5 cm/s, respectively. Neither in the first nor the second half of the night did we find significant differences between the median times or the median swim speeds of the three species (Mann-Whitney U-tests with Holm-Bonferroni correction, *p* > 0.05). Upstream movements were slower than downstream movements for all species (Mann-Whitney U-tests: *Sternopygus*: *u* = 686, *p* < 0.001, *Eigenmannia*: *u* = 222, *p* ≪ 0.001, *Apteronotus*: *u* = 9, *p* = 0.41), suggesting a distinct influence of the stream’s water flow on swim speed. From the median upstream and downstream swim speeds the water velocity Eq. (4) could be estimated to be about *v*_*stream*_ = 4.2 cm/s, which is at the low end of water velocities of lowland streams (Crampton, 1998). The average swim speeds Eq. (5) of the three species relative to the water can also be deduced: *Sternopygus*: *v*_*fish*_ = 22 cm/s, *Eigenmannia*: *v*_*fish*_ = 24 cm/s, *Apteronotus*: *v*_*fish*_ = 20 cm/s.

Based on the median speeds and times and assuming that the fish were continuously swimming upstream and then back downstream with constant speed and no interruptions by foraging, mating, etc., the maximum distances the fish could have traveled upstream within one night were 2.2 km, 3.0 km, and 1.1 km for *Sternopygus, Eigenmannia*, and *Apteronotus*, respectively. On the other hand, we found approximately 15 fish of each species hiding in root masses during the day per 100 m of stream, as measured along a 160 m long transect directly upstream of the electrode array. Thus, the number of fish we observed traversing the electrode array upstream in one night (Fig. 9 A) likely came from at least 330 m (*Sternopygus*), 180 m (*Eigenmannia*), and 50 m (*Apteronotus*) of stream adjoining our electrode array. However, as some fish appeared to be quite stationary within or close to our electrode array even at night (Fig. 5 A), traversing fish were likely recruited from longer stretches of the stream. Thus, the true distances covered by the moving fish in one night likely lie between several hundred meters and a few kilometers.

### EOD field characteristics

An important aspect of electric fish interactions are the effective EOD signal intensities at the position of a receiving fish. These are determined by the EOD amplitudes of the individual fish and their spatial distribution. The distribution of EOD potentials over the electrode array (Fig. 10 A, B) allowed us to infer the EOD amplitude and the exponent of the power-law decay *in situ* (Eq. 7; see methods).

For the example of a detailed measurement of a stationary *A. albifrons* recorded in a large outdoor tank (Fig. 2 A, Fig. 10 A), the estimated exponent *q* = 1.61 is similar to the one obtained by fitting the dipole potential based on Eq. (2) to the data (*q* = 1.63; Fig. 2 B). However, the estimated amplitude factor *P* = 24 mV cm^*q*^ was smaller than the one obtained from a dipole fit (*P* = 29 mV cm^*q*^). This bias to underestimate the amplitude results from estimating the maximum amplitude as an average over a substantial fraction of the data in each distance bin (here 5 %). Note, however, that for fish moving vertically offset from the electrode’s plane the proposed method profoundly underestimates both exponent and EOD amplitude. This effect is evident in the spatial EOD amplitude distribution and can be compensated by fitting the amplitude distribution over larger distances only. For the fish traversing the electrode array (Fig. 8 and Fig. 9), the exponents of the power-law decay of EOD amplitude with distance were on average *q* = 1.34±0.24 and did not differ significantly between the three species (Mann-Whitney U-tests, *p* > 0.25). The exponents are smaller than two because of boundary effects of the water surface and the stream bed (Fotowat et al., 2013).

The quantity that is measured by electroreceptor organs is the electric field, Eq. (8), i.e. the spatial derivative of the electric field potential (Fig. 10 C). Considering the known sensitivities of the electrosensory system for the closely related *A. albifrons* based on behavioral studies (0.5 µV/cm peak-to-peak ampltitude at 100 µS/cm, Knudsen, 1975), we estimated the effective detection range of *A. rostratus* from the field data according to Eq. (9) to 166±14 cm (*n* = 10) under the encountered natural conditions (Fig. 10 C). Similarly, we estimated the detection ranges for *Sternopygus* and *Eigenmannia* under the same conditions and using the same sensitivity (Fleishman et al., 1992; Knudsen, 1974, 1975) to be 255±76 cm (*n* = 97) and 253±54 cm (*n* = 64), respectively.

Our approach allows to compare EOD amplitudes within and across species. Usually, EOD amplitude is measured between a pair of electrodes positioned at the head and the tail of the fish. Since we did not know how large the fish were, we used the EOD amplitude at a distance of 50 cm computed from Eq. (7) as a robust measure for each fish’s EOD amplitude. The calculated EOD amplitudes were broadly distributed within each of the three species of wave-type weakly electric fish (*Sternopygus*: 0.47±0.30 mV; *Eigenmannia*: 0.46±0.23 mV; *Apteronotus*: 0.16±0.04 mV, Fig. 10 D). Whereas *Sternopygus* and *Eigenmannia* had similar EOD amplitude distributions (Mann-Whitney U-test *p* > 0.31), *Apteronotus* EOD amplitudes were clearly smaller than the former (Mann-Whitney U-test *p* ≪ 0.001). Only *Eigenmannia* showed a negative correlation between EOD amplitude and EOD *f* (Pearson’s *r* = −0.28 and *p* = 0.03, *Sternopygus*: *r* = 0.14, *p* = 0.17; *Apteronotus*: *r* = −0.13, *p* = 0.72).

**Figure 10:**
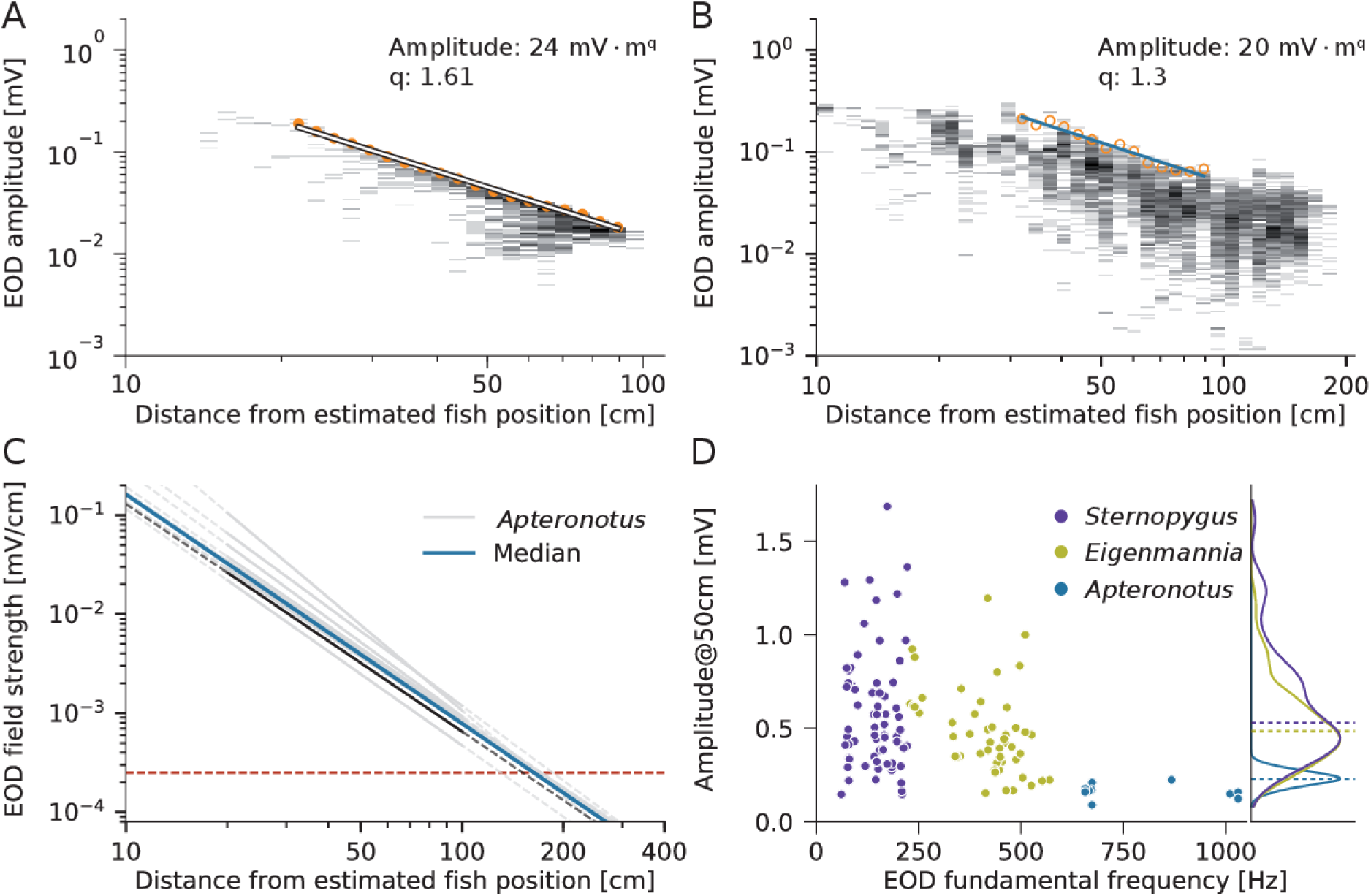
Estimation of EOD amplitude and its decay over distance. A) 2D-histogram of measures of EOD amplitudes obtained from all electrodes at various distances from the *A. albifrons* measured under controlled conditions in a tank (Fig. 2 A). Orange markers represent the mean of the largest 20% of EOD amplitudes per distance bin and were fitted with a power law function, Eq. (7) (line). B) 2D-histogram of EOD amplitudes of an example *A. rostratus* male (EOD*f* = 1030 Hz) measured from all electrodes at various distances. The means of the largest 5% of EOD amplitudes per distance bin (orange circles) were used for the estimate (blue) of the power-law decay, Eq. (7). C) EOD field strength, the gradient of the EOD amplitude, for the ten *A. rostratus* from Fig. 9 (gray lines; black: estimate from B, median in blue) with behavioral detection threshold (red horizontal dashed line; Knudsen, 1975). D) EOD amplitude estimates versus corresponding EOD frequencies for all three species. Histograms to the right show corresponding distributions of the EOD amplitudes, dashed lines indicate medians. *Sternopygus*: *n* = 69, *Eigenmannia*: *n* = 49, *Apternonotus*: *n* = 10.

## Discussion

We developed algorithms for tracking undisturbed and untagged individual wave-type electric fish based on their own, continuously active EOD. Using an electrode array we tracked the movements of three species of gymnotiform electric fish in their natural habitat during breeding season and characterized electric field properties of individual fish. We quantified nocturnal activity and revealed distinct movement patterns.

Previously, observing activity of wave- and pulse-type electric fishes in their natural habitats was based on transects recorded with single electrodes (e.g., Steinbach, 1970; Friedman and Hopkins, 1996). Small electrode arrays have so far been used only in controlled conditions in the lab (Jun et al., 2013; Matias et al., 2015) and for brief recordings in the field (Madhav et al., 2018). In our present study and in Henninger et al. (2018), we scaled this approach up to obtain data on natural behaviors of freely moving weakly electric fish in the field over extended periods of time.

### EOD-based individual tracking

Video tracking of behaving animals is well established for laboratory setups and allows for high throughput screening (Anderson and Perona, 2014; Mathis et al., 2018). Yet, video-based methods commonly rely on high contrast images and unobstructed line-of-sight, which make them challenging to use in natural habitats (Dell et al., 2014). Recent advances in miniaturization make loggers and transmitters attached to the animals an interesting opportunity for studying behavior in the wild (Cvikel et al., 2014; Krause et al., 2013; Strandburg-Peshkin et al., 2015; Flack et al., 2018). However, this approach requires to catch and recatch the animals in order to attach the logger and to retrieve the data. Importantly, it has been shown that tagging can have a negative impact on the animal’s fitness (Saraux et al., 2011). On the other hand, tagging induced only a transient stress response of cortisol levels, metabolic rate, and growth in killifish (Reemeyer et al., 2018).

In contrast, EOD-based tracking exclusively relies on the fish’s own, continuously emitted signals and therefore does not disturb the animals. This tracking approach allows for following individuals over time (Madhav et al., 2018), provides direct access to the fish’s communication signals (Henninger et al., 2018), and excels in terms of sub-second temporal resolution and a decent spatial resolution of about 10 cm. However, physical interactions like mouth wrestling or tail nipping, which could easily be detected visually in the lab (Triefenbach and Zakon, 2008), are inaccessible and can only be inferred indirectly, if at all, from EOD signals, context, and movement dynamics.

### Far-field characteristics of EODs

The electric near-field of electric fish is asymmetrically distorted along the body axis because of the elongated electric organ (Heiligenberg, 1973; Rasnow and Bower, 1996; Assad et al., 1998, 1999; Stoddard et al., 1999; Chen et al., 2005). At distances larger than about two body lengths the electric far-field approaches the one of an ideal dipole (Knudsen, 1975), independently of any asymmetries in the near field. Although the shape of the measured far-field can be well described by an ideal dipole model (Fig. 2 B), the exponent describing the decay of the field amplitude with distance is, with 1.6, clearly smaller than the exponent of 2 of the ideal dipole. The non-conducting bottom of the tank and the water surface induce boundary effects that compress the electric field (Fotowat et al., 2013) and result in a reduced exponent or even an exponential instead of power-law decay (Yu et al., 2019).

### EOD-based localization of fish

We computed estimates of the position of the fish based on the power at a fish’s EOD frequency measured on many electrodes of the array. Fitting a dipole model has been used successfully for estimating fish position (Jun et al., 2013; Madhav et al., 2018). However, these studies have not taken the reduced exponent in shallow water into account and assumed movements of the fish in the plane of the electrode array only. Our simulations showed that the dipole fit works well as long as the fish is in the grid’s center. However, the performance of the dipole fit deteriorates the farther the fish is above the plane of the electrode array and the closer it is at its borders (Fig. 3). A simple weighted average of the positions of the four electrodes with the largest signal amplitude turned out to be far more reliable, yielding localization errors below 10 cm, similar to the performance of the methods used in Madhav et al. (2018). In real-world scenarios, the dipole-like electric field can also be heavily distorted by objects like rocks, plants, and roots, where the fish like to hide. As long as such environments are not taken into account, model-based position estimates will be less precise and robust as well as computationally more demanding.

Note also, that we tested localization performance with *A. albifrons*, a low-amplitude species with an asymmetric near field (Fig. 2, Rasnow and Bower, 1996; Hoshimiya et al., 1980), and with simulations of an ideal dipole (Fig. 3). The overall dipole field, Eq. (2), is only multiplied by the EOD amplitude of a specific fish and by the water conductivity (Knudsen, 1975), we therefore do not expect profound differences in localization performance for species differing in EOD amplitude or lower conductivities. Localization performance was also independent of the degree of asymmetry of the near field (compare Fig. 2 C with Fig. 3 A).

Depending on the required spatial resolution, on the available electrode spacing, and on water conductivity/EOD amplitude, the water depth that can be covered by a 2D planar electrode array is limited. For the scenarios described here a planar electrode array covered a water depth of about 60 cm sufficiently well. The proposed methods can, however, easily be applied to electrode arrays arranged in multiple layers, because the weighted average Eq. (1) is directly based on the 3D corrdinates of the electrodes.

### Species-specific EOD frequencies

At our field site in Darién, Panamá, we found three syntopic wave-type species with quite well separated EOD *f* ranges: two Sternopygidae, *Sternopygus dariensis* with EOD frequencies below 220 Hz and *Eigenmannia humboldtii* at 200 – 580 Hz, and one member of the Apteronotidae, *Apteronotus rostratus*, with EOD*f* s ranging from 580 Hz to 1100 Hz. A similar separation between *Sternopygus* and *Eigenmannia* was previously found in Guyana (Hopkins, 1974b), coastal Surinam (Hopkins and Heiligenberg, 1978), Rio Negro (Bullock, 1969; Steinbach, 1970), and Napo River in eastern Ecuador (Stamper et al., 2010). EOD frequencies above 600 Hz are usually occupied by several sympatric species of gymnotiform fish. In Guyana *Apteronotus albifrons* overlapped with *Sternarchorhamphus macrostomus* (Hopkins, 1974a), in Rio Negro, five species shared frequencies above 800 Hz (Bullock, 1969; Steinbach, 1970), and in a whitewater river close to Manaus Kramer et al. (1981) found 28 species with EOD frequencies ranging from 300 to 1800 Hz.

Species identification based on EOD frequency was possible for our data, because only three species of wave-type fish are known for Panamá (Alda et al., 2013). The high densities of sympatric species sharing the same frequency band reported from the Amazon basin (e.g., Steinbach, 1970; Kramer et al., 1981) would pose a major challenge for future studies with an electrode array in these habitats. One would need to take into account additional characteristics of the EOD waveforms or the relative power of higher harmonics (Kramer et al., 1981; Turner et al., 2007).

Recognizing species that overlap in fundamental EOD frequency is not only a technical problem. Whether and how gymnotiform fish solve this problem themselves is not understood yet. *Eigenmannia* are able to discriminate female and male EOD waveforms even if they do not differ in EOD *f* (Kramer, 1999), and *Apteronotus leptorhynchus* were shown to chirp more to the signal of a real fish than to a sinewave mimic (Hopkins, 1974a; Dunlap and Larkins-Ford, 2003). However, in another playback experiment, signals based on the EOD waveforms of different species that overlapped in EOD frequencies failed to elicit differences in chirp responses in male *A. leptorhynchus*, although waveforms contained species-specific information (Fugère and Krahe, 2010).

Weakly electric fish do not need to rely on their baseline EOD alone. Species-specific modulations of the EOD frequency and amplitude could also be used to infer species identity (Kramer et al., 1981). Whereas rises seem to be highly conserved between species, chirps and EOD waveforms appear to be evolutionarily labile (Turner et al., 2007), and thus chirps are potential additional cues for species identification (Fugère and Krahe, 2010). The different types of chirps and EOD waveforms result in various degrees of conspicuousness of the resulting signals (Petzold et al., 2016) that may translate into discriminability by the electrosensory system. Chirps have also been suggested to aid disambiguation of the sign of high-frequency beats (Walz et al., 2014).

Note also, that the above mentioned field studies do not allow to resolve whether species are syntopic or cluster in separate microhabitats or in time (Kramer et al., 1981). Recordings of electric activity directly in the field (Stamper et al., 2010, and our data) clearly demonstrate that all three wave-type electric fish are both spatially and temporally coexisting in specific microhabitats.

### EOD interactions across species

Electroreceptor neurons are tuned to EOD frequencies and are most sensitive approximately at the EOD frequency of the individual fish (Hopkins, 1976). Behavioral thresholds in a detection task were similarly tuned (Knudsen, 1974). In playback experiments *Eigenmannia* responded to stimulus frequencies mimicking conspecifics (Hopkins, 1974a) and in *Sternopygus* with sexually dimorphic EOD frequencies males only responded to stimulus frequencies mimicking females (Hopkins, 1972, 1974b). During interactions of two electric fish, the difference between the two EOD frequencies is the relevant frequency for the tuning of P-type electroreceptor afferents (Bastian, 1981; Walz et al., 2014). P-type afferents thus encode relative and not absolute EOD frequency in their firing rate (but see Sinz et al., 2017). The question arises whether allospecific EOD frequencies are encoded by P-type afferents or whether electric fish have species-specific frequency channels?

Weakly electric fish are clearly able to sense the EODs of their conspecifics (Henninger et al., 2018). Since the frequency range covered by wave-type gymnotiform fish is rather broad (about half an octave, Hopkins, 1974b, Fig. 5 B), the frequency differences resulting from interacting conspecifics can be relatively large (Fig. 7 D). An impressive example is the courting behavior of a 1035 Hz male *Apteronotus rostratus* towards a 620 Hz female, resulting in an 415 Hz frequency difference (Henninger et al., 2018). However, although conspecifics may share a unique range of EOD frequencies, frequency differences do not allow to unambiguously distinguish species (Fig. 7 D). This ambiguity of frequency differences could be resolved if individual fish “knew” about their relative EOD frequency within their species’ EOD frequency range. Interestingly, electroreceptors of both *Eigenmannia* and *Sternopygus* are indeed more sharply tuned the higher their best frequency (Viancour, 1979; Zakon and Meyer, 1983). Whether this sharper tuning at the upper end of a species’ frequency range is enough to sufficiently suppress EODs of other species with higher EOD frequencies still needs to be shown. The related problem at the lower end of a species’ frequency range remains.

In addition, the harmonics of species with lower fundamental EOD frequencies fall into the sensitive range of the receptor tuning of a species with higher fundamental frequencies (Fig. 7 C). Although smaller in power, these harmonics might then create a beat that can be encoded by the receptor neurons. This should be a problem in *Apteronotus* species, in particular. Fish of this genus have much lower EOD amplitudes than *Sternopygus* and *Eigenmannia* (Fig. 10 D). Consequently, the harmonics of even a low-frequency *Sternopygus* can be of similar power to that of the fundamental frequency of *Apteronotus* (Fig. 6). This suggests that high-frequency fish should be able to perceive and interact with species of lower EOD frequencies. Indeed, *A. leptorhynchus* has been shown to increase chirp rate in response to playbacks and interactions with *Sternopygus macrurus* and *Brachyhypopomus gauderio* in the lab (Dunlap et al., 2010). Heterospecific interactions have also been reported for mormyrid electric fish (Scheffel and Kramer, 2006).

### Movement patterns

Our data from the electrode array allowed us to look beyond the mere presence of weakly electric fish species. Analyzing the movements of the recorded fish we found that about one third of the detections were fish that traversed the electrode array with or against the water flow (Fig. 8). At the beginning of their activity phase at the onset of the night fish of all three species were swimming upstream, whereas during the second half of the night the fish were preferentially swimming downstream (Fig. 9 A).

In contrast, a study on radio-tagged Australian graylings mainly found downstream movements over hundreds of meters and some upstream movements at the end of the night (Dawson and Koster, 2018). Based on transect data of general electric activity, Steinbach (1970) reported diurnal movements of gymnotiform fish over about one hundred meters from hiding places in deep waters during the day to shallow waters at night. Most available movement studies on fish in rivers, however, have not resolved diurnal movement patterns. Golden perch, carp, as well as the pulse-type weakly electric fish *Brachyhypopomus occidentalis*, for example, stayed within about one hundred meters for a few months, and only occasionally moved to a new home range farther away (Crook, 2004; Hagedorn, 1988).

We observed many upstream movements within and close to the root masses at the undercut bank, where water velocity was slowed down and where macroinvertebrates, potential prey, might hide and drift downstream. Downstream movements of the fish occurred more towards the shallower slip-off slope at the inner side of the stream’s bend with reduced water speeds. Based on the measured swimming speeds and on transect data we inferred a potential swimming range per night of at least a few hundreds meters and up to 3 km. This would be a rather large range for fish of about 20 cm length (Minns, 1995), but, as discussed above, most studies on fish migration behavior consider only movements from day to day, not within a single day. Depending on the taxon, movement distances can vary dramatically, but are positively correlated to fish length as well as stream size (Minns, 1995; Radinger and Wolter, 2014).

### Solitary fish

*Eigenmannia* has been reported as a social fish species that is usually found in groups (Hopkins, 1974a; Hagedorn and Heiligenberg, 1985). Tan et al. (2005) analyzed hundreds of snapshots of electric activity taken during the day and night in Ecuador. In most recordings multiple *Eigenmannias* were detected, but 10 to 20 % of the recordings contained only a single fish. In contrast, *Apteronotus* and *Sternopygus* have been mostly found solitary or in groups of two (Stamper et al., 2010). In the lab, groups of two are more frequent (Stamper et al., 2010), *Apteronotus* shows aggressive physical and electrocommunicative behavior (Triefenbach and Zakon, 2008), and only dominant males with the highest EOD frequency stay alone (Dunlap and Oliveri, 2002). In the field, however, aggressive electrocommunication seemed to be rare in *Apteronotus* (Henninger et al., 2018).

The analysis of our field data revealed that many fish were traversing the electrode array solitarily. This observation is in particular interesting for *Eigenmannia* and its jamming avoidance response (Heiligenberg, 1973; Behrend, 1977). Of the 65 traversing *Eigenmannia humboldtii* observed in a single night, 61 were swimming solitarily, a large number considering typical shoal sizes of about five that have been reported for *Eigenmannia virescens* (Tan et al., 2005). This suggests that a substantial number of fish disperse from their group during the night either to forage on their own or to find mating partners. Such nocturnal dispersion has been observed in the lab (Oestreich and Zakon, 2005). However, Tan et al. (2005) reported about similar numbers of *Eigenmannia* recorded on an electrode during the day and the night. Whether *Eigenmannia* forages solitarily or in shoals may also depend on the specific habitat and/or the availability of food, as has been shown for wild dogs (Hubel et al., 2016).

### Detection ranges

The data of the electrode array allowed us to characterize the spatial properties of the electric fields for individual fish *in situ* (Fig. 10 A) without any special recording procedure (Knudsen, 1975; Stoddard et al., 2007). For the given situation in the habitat, characterized by a certain water conductivity, electric field compression by the limited water column, and EOD strength, this allowed us to estimate the distance at which the electric field decayed down to the known behavioral sensitivities of several gymnotiform species to approximately 2 m. Within this range fish can detect and perceive each other, as we have shown in the field for attacks of a resident *A. rostratus* male towards an intruder (Henninger et al., 2018). The boundary effects of the water surface and the bottom, that effectively reduces the exponent of the power law decay (here *q* = 1.3) of the dipolar electric field (Fotowat et al., 2013; Yu et al., 2019), increase the detection ranges compared to a situation without boundaries and an exponent of two (Knudsen, 1975).

This detection range for conspecifics is much larger than typical distances of less than a body length for object detection (Knudsen, 1975). The minute changes induced by nearby objects are much smaller than modulations induced by EODs of con- and allospecific electric fish (Nelson and MacIver, 1999; Chen et al., 2005; Fotowat et al., 2013). By turning and bending the body the field strength and thus detectability at a receiving fish are further reduced even at small distances (Yu et al., 2019). The situation weakly electric fish face resembles walking with a dim flashlight through a forest in the dark: only the immediate environment can be illuminated by the flashlight, but the flashlight of another person can sometimes be seen over a much larger distance.

### EOD amplitudes

Our *in situ* characterization of electric fields also allowed us to estimate the amplitude of the electric organ discharges (Fig. 10 D). EOD amplitude is known to be strongly correlated with fish size (length or weight) within *Eigenmannia* (Westby and Kirschbaum, 1981), *Sternopygus* (Hopkins, 1972), *Brachyhypopomus* (Hagedorn, 1988), and also across species, including *Apteronotus albifrons* (Knudsen, 1975). Thus, the more than tenfold variation of EOD amplitudes that we estimated for *Sternopygus* and *Eigenmannia* seems to reflect considerable variability in fish sizes. The EOD amplitudes of the relatively few *Apteronotus* that we detected were much more homogeneous than those of the two other species. Most of them were involved in courtship behaviors and thus may have been above some minimum size (Henninger et al., 2018).

EOD frequency has also been reported to correlate with fish size and to signal dominance, for *Apteronotus leptorhynchus* in the lab (Hagedorn and Heiligenberg, 1985; Dunlap, 2002) and for *Sternarchorhynchus* in the wild (Fugère et al., 2011). In contrast, our data do not show any correlation between EOD frequency and EOD amplitude for *Sternopygus* and *Apteronotus* (Fig. 10 D).

### Natural sensory scenes

By far most of our knowledge about the behavior of weakly electric fish is based on studies of captive fish in tanks. Quantitative observation of any fish species in their natural habitats has been next to impossible without tagging, even more so in species that exhibit a nocturnal and hidden life style. In contrast to non-electric fish, weakly electric fish can be detected based alone on their continuous electric organ discharges. This has been exploited in many of the outdoor studies, mostly by means of single electrodes used to locate the fish (e.g., Bullock, 1969; Steinbach, 1970; Hopkins, 1974a; Kramer et al., 1981; Hagedorn, 1988; Westby, 1988; Friedman and Hopkins, 1996; Tan et al., 2005; Stamper et al., 2010). Expanding this technique to arrays of tens of electrodes that monitor the activity of weakly electric fish over a whole day yields a plethora of valuable information on the secret lives of weakly electric fishes in tropical habitats and the associated natural electrosensory scenes that need to be processed by their electrosensory system. In Henninger et al. (2018), we found in courtship and aggression contexts of *Apteronotus* behaviorally relevant electrical signals of unexpected frequencies and amplitudes that have so far been largely neglected in neurophysiological studies.

The multispecies community we describe here (Fig. 5 and Fig. 7) hints at the complexity of signals gymnotiform fish are actually facing. Most of the behavioral and electrophysiological literature so far has focused on static interactions of two fish, but already relative movements induce higher order amplitude modulations of the resulting signals, so called “envelopes” (Yu et al., 2012, 2019). The interaction of more than two fish also results in envelopes and in specific behavioral responses (Partridge and Heiligenberg, 1980; Stamper et al., 2012). Courtship and agression behaviors of *Apteronotus* in our recordings demonstrate that the fish are able to selectively respond to specific fish in the presence of other nearby fish (Henninger et al., 2018). Here we introduced algorithms that allow to extract EOD frequencies and amplitudes of individual fish as well as distances between interacting fish, providing the basis for a detailed quantification of the complexity of social signals in natural scenes in the future.

The movements of the fish traversing the electrode array (Fig. 8) in combination with transect data along the stream hint at long-range navigation abilities of the fish in the range of several hundred meters up to potentially a few kilometers. In small-scale laboratory settings, gymnotiforms can be trained in spatial orientation tasks (Jun et al., 2016) and hippocampal-like circuitry has been described in their pallium (Elliott et al., 2017). How far the fish really travel along the stream, whether they return back to some preferred hiding place, and what cues they rely on (electrical, lateral line, visual, or olfactory) are exciting questions for future studies with multiple smaller electrode arrays distributed along a stream.

## Acknowledgements

We thank Hans Reiner Polder and Jürgen Planck from npi electronic GmbH for designing the amplifier, Sophie Picq, Diana Sharpe, Luis de León Reyna, Rigoberto González, Eldredge Bermingham, the staff from the Smithsonian Tropical Research Institute, and the Emberá community of Peña Bijagual for their logistical support.

